# Greater distal activation of the biceps femoris long head reflects proximodistal differences in motor unit action potential properties

**DOI:** 10.1101/2024.10.14.618251

**Authors:** José Carlos dos Santos Albarello, Hélio V. Cabral, Francesco Negro, Liliam Fernandes de Oliveira

**Author notes:** <u>Corresponding authors:</u> Dr. Hélio V. Cabral, Department of Clinical and Experimental Sciences Università degli Studi di Brescia, Viale Europa 11, Brescia, 25121, Italy. These authors contributed equally to this work and shared senior authorship (Francesco Negro, Liliam Fernandes de Oliveira).

## Abstract

**Purpose:** Recent research has explored region-specific responses within the biceps femoris long head. However, evidence on regional muscle activation remains controversial, primarily because information derived solely from surface electromyograms (sEMG) amplitude does not necessarily provide an accurate estimate of neural drive to the muscle. To address this limitation, this study investigated whether there are proximodistal differences in motor unit properties of the biceps femoris long head during isometric hip extension and knee flexion tasks.

**Methods:** Seventeen resistance-trained males performed isometric knee flexion and hip extension tasks at 20% and 40% of maximal voluntary contraction. High-density sEMG were recorded from proximal and distal regions of the biceps femoris long head and decomposed into individual motor units. Central motor unit properties (mean discharge rate, discharge rate variability, recruitment and de-recruitment thresholds) and action potential properties (amplitude and conduction velocity) were analyzed. Bipolar sEMG amplitude was also calculated for each region to simulate traditional sEMG measurements.

**Results:** Bipolar sEMG amplitude, motor unit action potential amplitude and conduction velocity were significantly greater in the distal region during both tasks. In contrast, no proximodistal differences were observed in central motor unit properties.

**Conclusion:** These findings suggest that increased bipolar sEMG amplitude in the distal region of the biceps femoris long head is driven by motor unit action potential properties rather than differences in central modulation, likely influenced by intra-muscular variations in muscle mechanics and geometry. This emphasizes limitations of relying solely on sEMG amplitude to infer neural control strategies in the biceps femoris long head.

## Introduction

Predominantly composed of biarticular muscles (apart from the biceps femoris short head), the hamstrings play a key role in tasks requiring hip extension and knee flexion movements and are therefore engaged in many daily living activities (Afonso et al. 2021; Kellis and Blazevich 2022). For instance, during the leg swing phase in running or sprinting, the hamstrings contract eccentrically to decelerate the hip flexion and knee extension, thereby greatly contributing to the stabilization of these joints during directional changes (Fields et al. 2005; Hogervorst and Vereecke 2015). However, the substantial eccentric load at the end of the swing phase, when the hamstrings are at longer lengths, makes this muscle group highly susceptible to stretch-induced injuries, particularly during high-velocity movements (Jonhagen et al.; Liu et al. 2012; Danielsson et al. 2020; Jokela et al. 2023). Indeed, the incidence of hamstring injuries in sports has systematically increased over the last years (Ekstrand et al. 2023), with still high prevalence rates (Zachazewski et al. 2019; Ribeiro-Alvares et al. 2020). Despite significant efforts by the scientific community to understand the underlying mechanisms and develop strategies for prevention and treatment (Brukner 2015), the rate of hamstring injuries has remained relatively unchanged over the past 30 years (Maniar et al. 2022). This persistent trend underscores the need for further research to address existing gaps in this literature, aiming to reduce the risk of hamstrings injuries. Of particular interest, a wide body of research has recently explored region-specific responses within the hamstring muscles, particularly in the proximal and distal regions of the biceps femoris long head (Schoenfeld et al. 2015; Watanabe et al. 2016; Hegyi et al. 2019), which is the most frequently injured muscle among the hamstrings (Opar et al., 2012). The possibility of a proximodistal selective response in the biceps femoris long head during hip extension and knee flexion tasks primarily arises from existing evidence suggesting task-dependent regional activation in biarticular muscles (Watanabe et al. 2021), such as the rectus femoris (Miyamoto et al. 2012; Watanabe et al. 2012) and the gastrocnemius (Wolf et al. 1993; Cohen et al. 2020) muscles. However, evidence regarding the biceps femoris long head muscle remains controversial. For instance, Watanabe et al. (2016) observed no differences in the proximodistal activation of the biceps femoris long head muscle during knee flexion and hip extension isometric tasks. Conversely, Hegyi et al. (2019) reported greater activation in the distal region compared to the proximal region during the hamstring curl exercise (which exclusively involves knee flexion) but found no proximodistal differences during hip extension exercises. These inconsistent findings may be attributed to the fact that information extracted solely from the amplitude of surface electromyograms (sEMG) does not necessarily provide an accurate estimate of the neural drive to the muscle due to a phenomenon known as “amplitude cancellation” (Keenan et al. 2005; Farina et al. 2010). Moreover, sEMG amplitude is also influenced by various peripheral factors, such as muscle architecture and motor unit conduction velocity (Farina et al. 2004, 2014), further complicating the interpretation of changes in sEMG amplitude as changes in the neural drive to the muscle (Martinez-Valdes et al. 2018).

These limitations can be overcome by combining blind source separation methods with high-density surface electromyogram (HDsEMG) recordings, which allow for the noninvasive decomposition and assessment of the activity of individual motor units (Holobar and Zazula 2007; Negro et al. 2016). These methods enable the direct study of central and peripheral properties of motor units located in different muscles or muscle regions (Martinez-Valdes et al. 2018; Valli et al. 2024), providing a more precise estimate of potential proximodistal differences in the control of the biceps femoris long head muscle. Therefore, in this study, we investigated whether there are proximodistal differences in the motor unit properties of the biceps femoris long head during hip extension and knee flexion isometric tasks. To achieve this, we decomposed motor unit spike trains from HDsEMG acquired from the proximal and distal regions of the biceps femoris long head and compared central (mean discharge rate, discharge rate variability, recruitment threshold and de-recruitment threshold) and action potential (motor unit conduction velocity and motor unit action potential amplitude) properties between regions. Additionally, to compare our findings with proximodistal differences in bipolar sEMG, as previously reported (Schoenfeld et al. 2015), we simulated two bipolar sEMG detection sites in the proximal and distal regions and calculated their amplitude, both normalized and non-normalized.

## Methods

### Participants

Seventeen healthy men (mean ± SD; age: 30 ± 4 years; height: 176.82 ± 6.09 cm; body mass: 84.71 ± 10.46 kg) participated in this study. All participants were resistance training practitioners and reported no lower limb musculoskeletal injuries in the preceding 6 months. Prior to the start of experiments, all participants read and signed an informed consent form. The study was conducted in accordance with the latest version of the Declaration of Helsinki and was approved by the ethics committee of the Clementino Fraga Filho University Hospital of the Federal University of Rio de Janeiro (approval number: 6.259.296).

### Experimental protocol

All experimental procedures were carried out in a single visit to the laboratory lasting approximately 90 minutes. Initially, participants were familiarized with the equipment and tasks, followed by measurements of body mass and height. Then, participants were asked to perform isometric torque-matching tasks involving knee flexion and hip extension contractions (**Figure 1A**). For the knee flexion task, participants lay prone on the dynamometer chair, with the knee joint axis aligned as coaxially as possible with the dynamometer’s axis of rotation (left panel in **Figure 1A**). For the hip extension task, participants stood on the shaft of the dynamometer, aligning the hip joint as coaxially as possible with the dynamometer’s axis of rotation (right panel in **Figure 1A**). These positions were chosen to maintain the biceps femoris at approximately the same muscle length between conditions.

**Figure 1:**
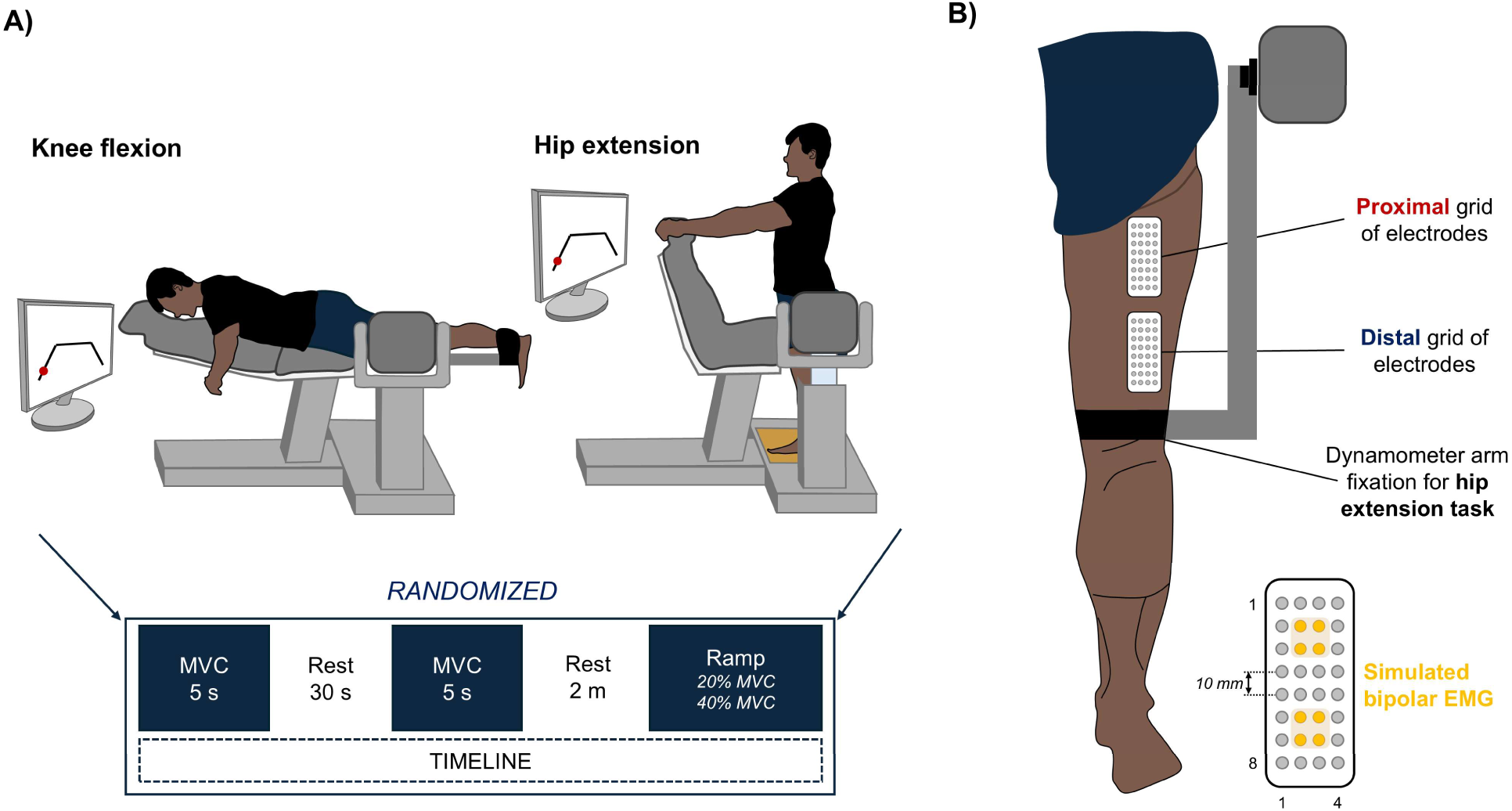
Experimental procedures. (A) Participants performed isometric knee flexion and hip extension tasks. Two maximal voluntary contractions (MVCs) were conducted for each task followed by trapezoidal ramp contractions were at 20% and 40% of MVC (bottom part). (B) High-density surface electromyograms were recorded from the proximal and distal regions of the biceps femoris long head using grids of 32 electrode each. The yellow-highlighted electrodes indicate the specific channels used to simulate a bipolar sEMG configuration.

Each task followed the same experimental protocol (bottom part of **Figure 1A**). First, participants performed two maximum voluntary isometric contractions (MVC) for 5 seconds, with a 30-second interval between contractions. The highest value from the two MVCs was considered for the subsequent submaximal isometric tasks. After a 2-minute rest, participants were asked to perform isometric trapezoidal ramps consisting of a linear increase from 0% MVC to the target torque level in 5 seconds, a plateau phase at the target torque level for 30 seconds, and a linear decrease from the target torque level to 0% MVC in 5 seconds. Considering that contraction intensity can affect the neural drive to active motor units (Martinez-Valdes et al. 2022), hip extension and knee flexion isometric tasks were performed with two different intensities. Consequently, participants performed four isometric torque-matching tasks in total: knee flexion at 20% MVC, knee flexion at 40% MVC, hip extension at 20% MVC, hip extension at 40% MVC. The order of the tasks (knee flexion and hip extension) was randomized.

### Data collection

During the tasks, two semi-disposable 32-channel grids of electrodes (10 mm interelectrode distance, GR10MM0804, OTBioelettronica, Turin, Italy; bottom part of Figure 1B) were used to record the activity of the long head of the biceps femoris muscle. Before electrode placement, the skin was shaved and cleaned with neutral soap and water. With participants lying prone on a stretcher, the electrodes were positioned on the proximal and distal regions of the long head of the biceps femoris (Figure 1B). Ultrasound imaging (Logiq, GE Healthcare, USA) was used to identify and mark the aponeuroses of the biceps femoris muscle on the skin, ensuring precise grid placement. The grids were positioned as proximally and distally as possible on the muscle, aligned longitudinally with the fascicles, and not over adjacent muscles. The HDsEMG signals were recorded in monopolar mode and digitalized synchronously with the torque output at a sampling frequency of 2000 Hz using a 16-bit wireless amplifier (10–500 Hz bandwidth; Gain 1; MUOVI, OTBioelettronica, Turin, Italy).

### Data analysis

HDsEMG signals were analyzed offline using MATLAB (version 2022b, The MathWorks Inc., Natick, Massachusetts, USA) custom-written scripts. Monopolar EMGs were digitally filtered between 20 and 500 Hz using a second-order Butterworth band-pass filter. Each monopolar signal was visually inspected, and channels deemed unsuitable for analysis (e.g., containing artifacts) were excluded.

#### Bipolar sEMG amplitude

To compare proximodistal differences in the bipolar sEMG amplitude of the long head of the biceps femoris during knee flexion and hip extension tasks, we simulated two bipolar sEMG detection sites: one in the proximal region and one in the distal region. For each grid, we calculated the difference between the average monopolar signals from groups of four channels in a proximal and a distal location within the electrode grid (yellow channels indicated in the bottom part of Figure 1B). To quantify the biceps femoris muscle activation, the root-mean-square (RMS) amplitude was calculated in non-overlapping windows of 250 ms during the 30-s plateau phase, and the average RMS value across these windows was considered for further analysis. Additionally, we calculated the normalized RMS amplitude by dividing the average RMS value by the RMS amplitude during the MVC, which was computed as the RMS value in a 500-ms time window centered around the peak torque. The maximum MVC RMS value between proximal and distal bipolar sEMG was considered for the normalization.

#### Motor unit decomposition

HDsEMG signals from the proximal and distal grids were decomposed separately into individual motor unit spike trains (Figure 2A) using a convolutive blind source separation method (Negro et al. 2016). Briefly, after performing the convolutive sphering of the HDsEMG signals (i.e., extending, centering, and whitening), a fixed-point algorithm was applied to identify sources (i.e., motor unit spike trains) that maximize a measure of sparsity. Spikes were separated from noise using K-means clustering, and as the motor unit separation vectors were iteratively updated, the estimation of discharge times was further refined by minimizing the CoV of the inter-spike intervals (for details, see Negro et al., 2016). Motor units automatically identified by the algorithm were individually inspected and, when necessary, manually edited by an experienced operator (Martinez-Valdes et al. 2023). The editing step is performed to improve the accuracy of the identified motor unit discharge times, primarily by removing misidentified discharges (e.g., inter-spike intervals < 20 ms) and adding missing dischargers (e.g., inter-spike intervals > 250 ms). To ensure the analyzed motor units were exclusively from the proximal and distal regions, duplicated motor units (i.e., those identified in both grids) were excluded from further analysis. To this end, motor unit spike trains from the proximal and distal regions were compared and considered to be generated by the same motor unit if they shared at least 30% of discharge times (Negro et al. 2016).

**Figure 2:**
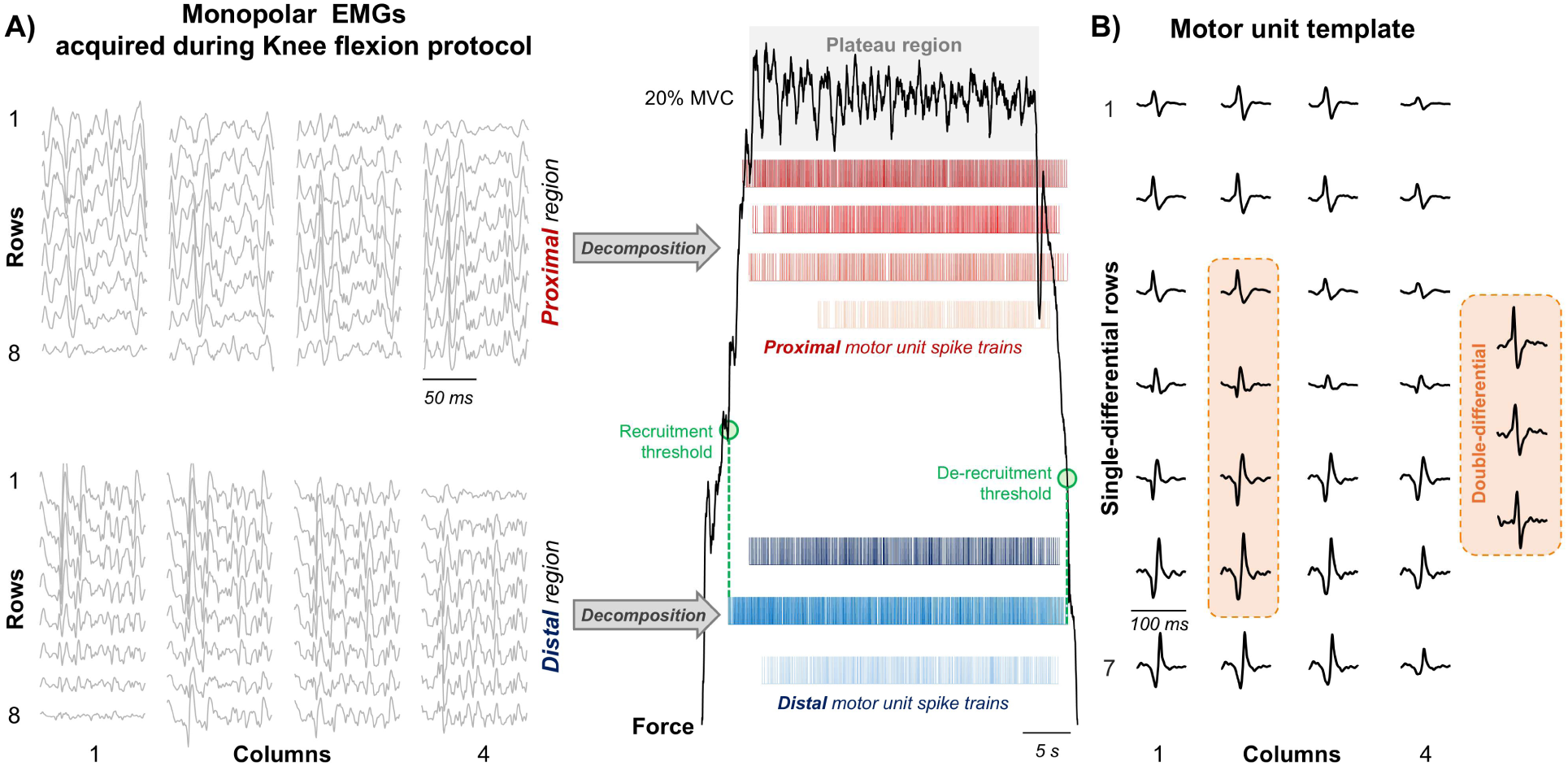
Calculation of motor unit properties. (A) Monopolar surface electromyograms (sEMG) were decomposed into individual motor unit spike trains for each muscle region. Recruitment and de-recruitment thresholds were identified as the torque level (% maximal voluntary contraction) at which the motor unit first began and stop to discharge (green dashed lines), respectively. The mean discharge rate and discharge rate variability were calculated from the plateau phase (light grey rectangle) of the torque output (black line). (B) Motor unit action potential templates were obtained by triggering and averaging the single-differential sEMG within a 50 ms window centered on motor unit discharge times. The motor unit action potentials’ peak-to-peak amplitudes were calculated from both the simulated bipolar sEMG detection sites and from the active channels of the grid. Motor unit conduction velocity was estimated using a multichannel maximum likelihood algorithm, selecting from three to six consecutive double-differential channels that displayed the clearest action potential propagation in each column of the grid (orange rectangle).

#### Motor unit central properties

To quantify proximodistal differences in motor unit central properties of the long head of the biceps femoris muscle during knee flexion and hip extension tasks, the mean discharge rate, the discharge rate variability, the recruitment threshold, and the de-recruitment threshold were calculated for each motor unit. The instantaneous discharge rate of each motor unit was calculated from the multiplicative inverse of the inter-spike interval and the mean discharge rate during the plateau phase was considered for analysis. The motor unit discharge rate variability was computed as the CoV of the inter-spike interval (CoV-ISI). The motor unit recruitment threshold and de-recruitment threshold were determined by identifying the torque level (%MVC) at which the motor unit first began and stop to discharge during the contraction, respectively (green-dashed vertical lines in Figure 2A) (Gomes et al. 2024).

#### Motor unit action potential properties

To quantify proximodistal differences in motor unit action potential (MUAP) properties of the long head of the biceps femoris muscle during knee flexion and hip extension tasks, the MUAP amplitude and conduction velocity were calculated. MUAP templates were estimated by triggering and averaging the single-differential sEMG signals at 50 ms windows centered at the discharge times of individual motor units (i.e., spike-triggered averaging technique; Stein et al. 1972; Figure 2B). Two indices were used to assess MUAP amplitude: (i) the peak-to-peak amplitude of the MUAP at the simulated bipolar sEMG detection site (see *Bipolar sEMG amplitude* section), and (ii) the average peak-to-peak amplitude across the MUAPs from the active channels of the grid. Active channels were defined as those with a peak-to-peak amplitude greater than 70% of the maximum amplitude observed across the grid (Vieira et al. 2010). The MUAP conduction velocity was measured from a minimum of three to a maximum of six double-differential channels in each column of the grid using the multichannel maximum likelihood algorithm (Farina et al. 2001). For each column, the channels that displayed the clearest action potentials propagation were manually selected (see orange rectangle in Figure 2B), and the conduction velocity estimated from the column with the higher cross-correlation across selected channels was considered for further analysis (Valli et al. 2024).

### Statistics

Linear mixed-effect models (LMM) were applied to compare motor unit central properties (mean discharge rate, CoV-ISI, recruitment threshold and de-recruitment threshold), motor unit action potential properties (MUAP amplitude and MUAP conduction velocity), and bipolar sEMG amplitude (raw and normalized) between the proximal and distal regions, and torque levels. For each task, random intercept models were used with region (proximal and distal) and torque level (20% and 40% MVC) as fixed effects and participant as random effect (*i.e., motor unit property or sEMG amplitude ∼ 1 + region*torque + (1 | participant*)). These models were implemented using the package *lmerTest* (Kuznetsova et al. 2017) with the Kenward-Roger method to approximate the degrees of freedom and estimate the *P*-value. Q-Q plot inspection was used to assess the assumption of normality of residuals. The *emmeans* package was used, when necessary, for multiple comparisons and to determine estimated marginal means with 95% confidence intervals (Lenth et al. 2019). All statistical analyzes were conducted with the R programming language (version 4.2.0). All individual data are provided as supplementary material.

## Results

### Proximodistal differences in bipolar sEMG amplitude

To investigate proximodistal differences in the biceps femoris long head activation during knee flexion and hip extension tasks, we simulated two bipolar sEMG detection sites in each region and calculated the RMS amplitude value. For both the raw and normalized RMS values and for all investigated tasks, significantly greater bipolar sEMG amplitudes were detected at the distal region compared to the proximal region (LMM; F > 9.87; *P* < 0.002 for all cases), and at 40% MVC compared to 20% MVC (LMM; F > 56.68; *P* < 0.001 for all cases). Additionally, no significant interactions between torque level and region were observed (LMM; F < 3.64; *P* > 0.06 for all cases). At 20% MVC, the raw RMS amplitude values (mean ± SD) for knee flexion were 42.9 ± 26.9 µV in the proximal region and 74.9 ± 37.8 µV in the distal region. For hip extension, the values were 58.1 ± 34.9 µV in the proximal region and 90.8 ± 72.9 µV in the distal region. At 40% MVC, knee flexion showed 139 ± 58.3 µV in the proximal region and 201 ± 55.5 µV in the distal region, while hip extension showed 131 ± 67.9 µV in the proximal region and 166 ± 84.7 µV in the distal region. Normalized bipolar RMS amplitude results are presented in Figure 3.

**Figure 3:**
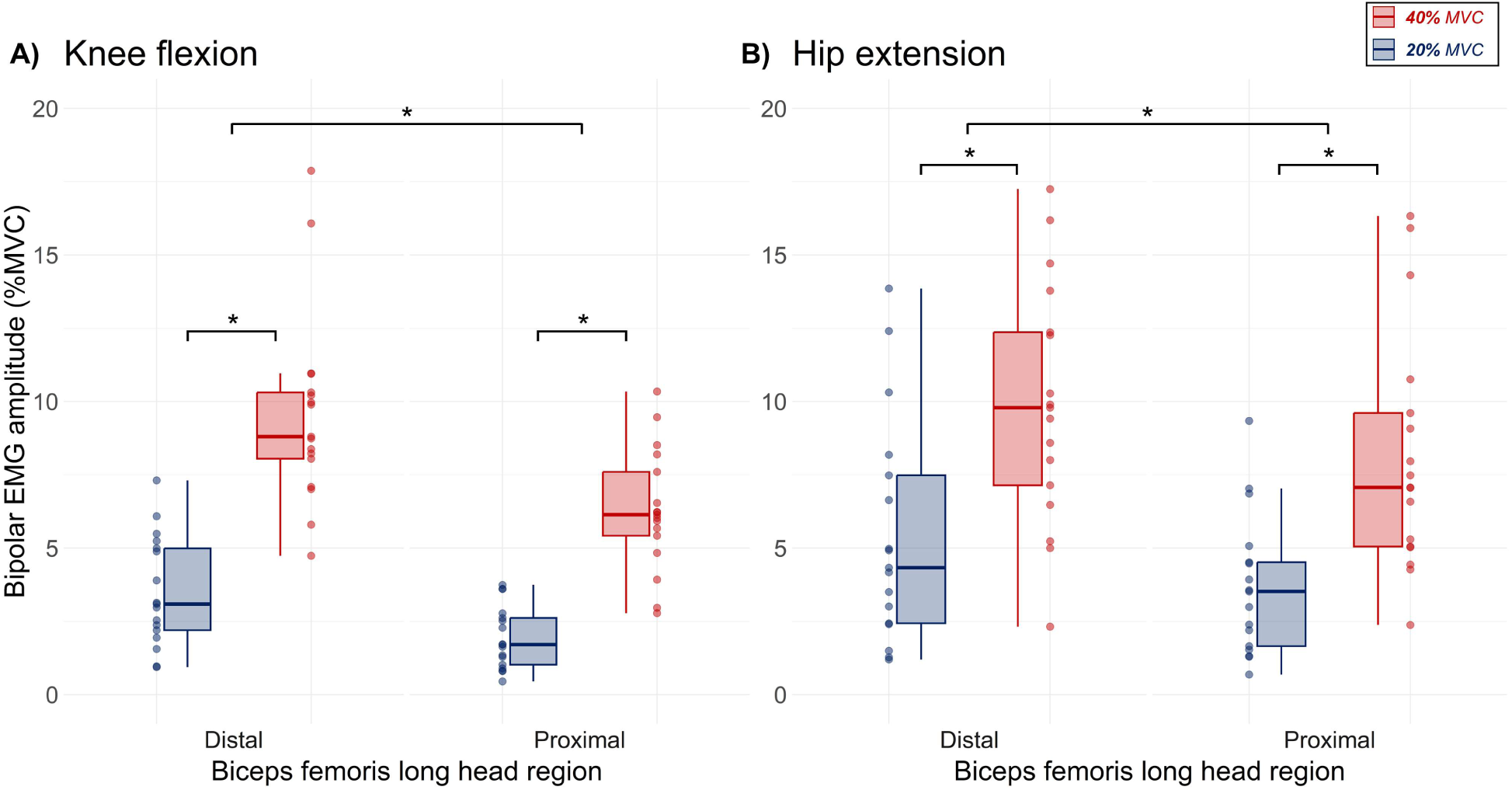
Bipolar EMG amplitude results. Normalized root-mean-square amplitude values from simulated bipolar sEMG at 20% of maximal voluntary contraction (MVC) (blue) and 40% MVC (red), separately for knee flexion (A) and hip extension (B) tasks. Horizontal traces, boxes, and whiskers, respectively, denote median value, interquartile interval, and distribution range. Asterisk denotes statistical differences between distal and proximal muscle regions (P < 0.05).

### Proximodistal differences in motor unit central properties

To assess whether there were proximodistal differences in motor unit central properties of the biceps femoris long head during knee flexion and hip extension tasks, we compared the mean discharge rate, discharge rate variability (CoV-ISI), recruitment threshold and de-recruitment threshold of decomposed motor units. When pooling motor units from the proximal and distal regions, an average of 6 ± 4 and 5 ± 4 motor units were identified per participant for the knee flexion at 20% MVC and knee flexion at 40% MVC, respectively. For hip extension tasks, 6 ± 5 and 3 ± 3 motor units were identified per participant for the hip extension at 20% MVC and hip extension at 40% MVC, respectively.

For all the tasks, no significant differences in motor unit mean discharge rate were observed between the proximal and distal regions of the biceps femoris long head (LMM; F < 1.20; *P* > 0.274 for all cases; Figure 4). Conversely, significant increases in mean discharge rate of approximately 1.8 pps and 4.1 pps were observed between 20% and 40% MVC for knee flexion (Figure 4A) and hip extension (Figure 4B), respectively (LMM; F > 14.37; *P* < 0.001 for all cases).

**Figure 4:**
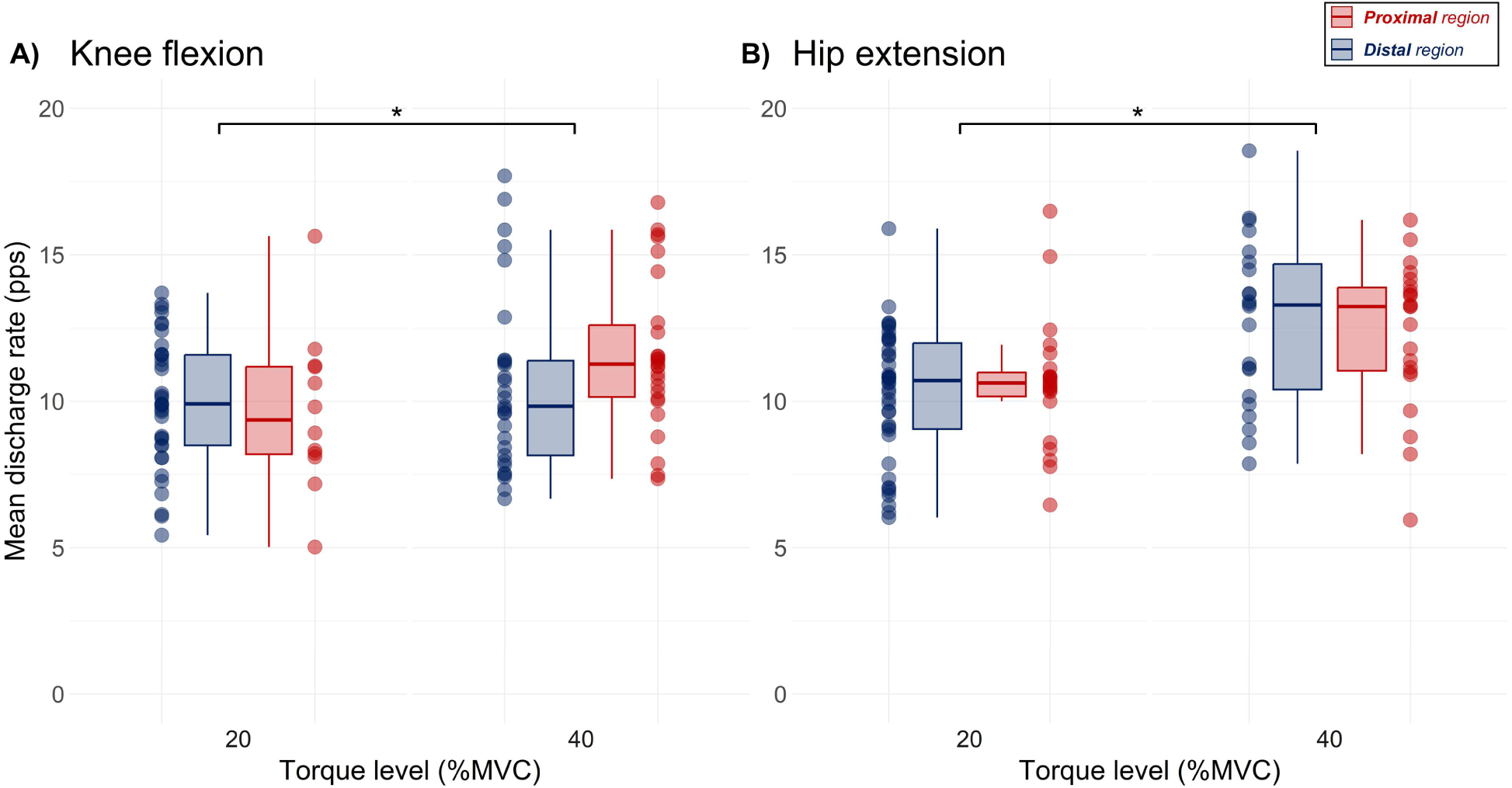
Mean discharge rate results. Mean discharge rate values of motor units decomposed from the proximal (red) and distal (blue) regions of the biceps femoris long head, separately for knee flexion (A) and hip extension (B) tasks. Data is grouped by torque level. Horizontal traces, boxes, and whiskers, respectively, denote median value, interquartile interval, and distribution range. Asterisk denotes statistical differences between 20% and 40% of maximal voluntary contraction (MVC) (P < 0.05).

Regarding motor unit discharge rate variability, significant increases of ∼3% in CoV-ISI were observed between 20% and 40% MVC only for the knee flexion task (LMM; F = 6.65; *P* = 0.012). No significant differences were observed across torque levels (LMM; F = 0.975; *P* = 0.326) and between the proximal and distal regions of the biceps femoris long head for the hip extension task (LMM; F = 2.23; *P* = 0.634). At 20% MVC, the CoV-ISI values (mean ± SD) for knee flexion were 16.5 ± 4.21% in the distal region and 15.9 ± 4.53% in the proximal region, while for hip extension the values were 19.3 ± 6.00% and 21.6 ± 7.80% in the distal and proximal regions respectively. At 40% MVC, knee flexion showed 22.2 ± 7.18% in the distal region and 19.8 ± 6.11% in the proximal, while hip extension showed 21.7 ± 6.20% and 21.0 ± 5.79% in the distal and proximal regions, respectively.

For both tasks, significant increases in the motor unit recruitment threshold (LMM; F > 133.56; *P* < 0.001 for all cases; **Online Resource 1**) and de-recruitment threshold (LMM; F > 244.89; *P* < 0.001 for all cases; **Online Resource 2**) were found between 20% and 40% of MVC. Conversely, no significant differences in recruitment and de-recruitment thresholds were observed between the proximal and distal regions (LMM; F < 2.058; *P* > 0.119 for all cases).

### Proximodistal differences in motor unit action potential properties

To evaluate proximodistal differences in MUAP properties, we calculated the MUAP amplitude as (i) the peak-to-peak amplitude of MUAP at the simulated bipolar sEMG site, and (ii) the average peak-to-peak amplitude across the MUAPs from the active channels of the grid. For the knee flexion and hip extension tasks and both MUAP amplitude indexes, significant great amplitudes were observed for the distal region of the biceps femoris long head (LMM; F > 4.04; *P* > 0.049 for all cases), except for the MUAP at the simulated bipolar sEMG during the knee flexion task (LMM; F = 3.54, P = 0.064). Additionally, significant increases in both MUAP amplitude indexes were observed between 20% and 40% MVC (LMM; F > 7.42; *P* < 0.008 for all cases). The MUAP amplitude from active channels are presented in Figure 5.

**Figure 5:**
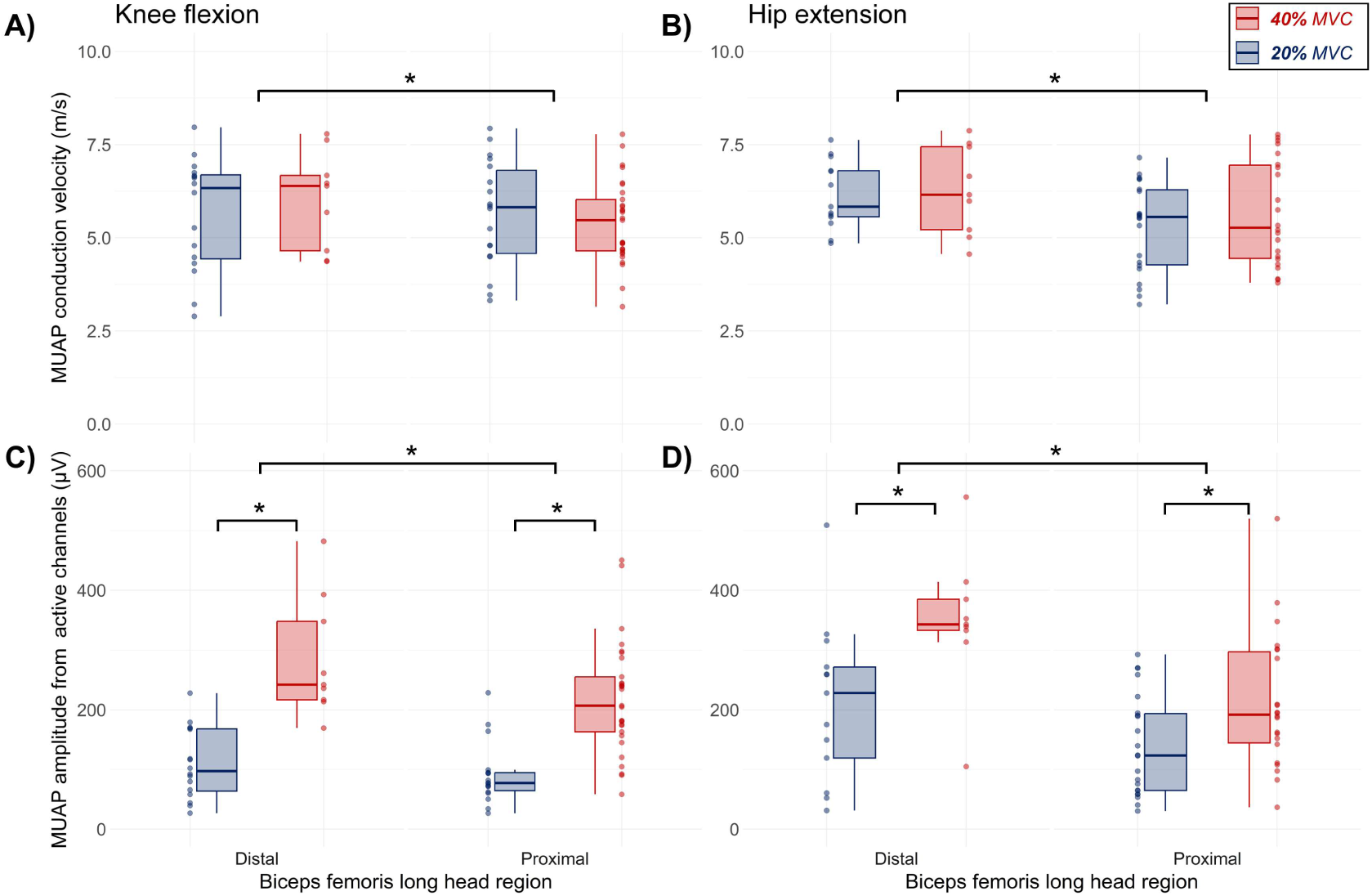
Motor unit action potential amplitude and conduction velocity results. Motor unit action potential amplitude from active channels and motor unit conduction velocity results of motor units decomposed at 20% of maximal voluntary contraction (MVC) (blue) and 40% MVC (red), separately for knee flexion (A and C) and hip extension (B and D) tasks. Horizontal traces, boxes, and whiskers, respectively, denote median value, interquartile interval, and distribution range. Asterisk denotes statistical differences between distal and proximal muscle regions (P < 0.05).

We also quantified MUAP conduction velocity to assess proximodistal differences in MUAP properties. For both knee flexion (Figure 5A) and hip extension (Figure 5B) tasks, significantly higher MUAP conduction velocities were observed in the distal region compared to the proximal region (LMM; F > 4.45; *P* < 0.038 for all cases). Additionally, no significant main effect of torque level or interactions between torque level and region were observed (LMM; F < 1.39; *P* > 0.241 for all cases).

## Discussion

The present study investigated proximodistal differences in the activity of the biceps femoris long head during isometric hip extension and knee flexion tasks. Our primary findings demonstrated consistently greater bipolar sEMG amplitudes at the distal region compared to the proximal region, regardless of the task performed. Additionally, while no significant differences were observed in the motor unit central properties, the distal region showed significantly higher MUAP conduction velocities and MUAP amplitudes than the proximal region. As discussed below, these results suggest that the increased activation in the distal region of the biceps femoris long head during both knee flexion and hip extension tasks is not related to differences in neural control of motor units between regions, but it is likely attributed to the size and characteristics of the motor unit action potentials.

### Proximodistal activation within the biceps femoris long head during knee flexion and hip extension tasks

Regional activation within human skeletal muscles has been consistently demonstrated during voluntary contractions, particularly in muscles that contribute to multiple joint actions (Staudenmann et al. 2009; Watanabe et al. 2012; de Souza et al. 2017; Cabral et al. 2022; Albarello et al. 2022). This non-uniform activation requires a specific neuroanatomical organization, wherein multiple branches of the nerve innervate localized regions within the muscle (i.e., motor unit with relatively small territories; Watanabe et al., 2021). Such patterns have been previously observed in the gastrocnemius (Segal et al. 1991) and rectus femoris (Yang and Morris 1999). Notably, the biceps femoris muscle is innervated by two branches of the sciatic nerve, which insert at multiple locations along its proximodistal axes (Seidel et al. 1996). Given this innervation pattern, it is reasonable to hypothesize region-specific activation within this muscle.

Consistent with this hypothesis, our study reported localized activation of the biceps femoris long head during both knee flexion and hip extension tasks, with the distal region exhibiting consistently higher bipolar sEMG amplitude compared to the proximal region (Figure 3). These results align with previous studies that reported greater sEMG amplitudes in the distal region during knee flexion tasks (Schoenfeld et al. 2015; Hegyi et al. 2018, 2019; Miyamoto and Hirata 2021). However, in contrast to our findings, previous evidence has suggested a more homogeneous proximodistal activation pattern within the biceps femoris during hip extension tasks (Schoenfeld et al. 2015; Hegyi et al. 2018). These discrepancies may be attributable to methodological differences, particularly in contraction type and task specificity. Our study used isometric tasks without torque at the knee joint (the tested limb was slightly suspended, and the dynamometer arm was fixed above the popliteal fossa; see Figure 1B), which may differ from the activation patterns in dynamic conditions. Additionally, methodological factors must be considered when interpreting sEMG amplitude as an indicator of neural drive to the muscle.

Ideally, the sEMG signal reflects the strength of the neural drive to the muscle and its amplitude properties have traditionally been used to infer neural control strategies (Del Vecchio et al. 2017). However, sEMG amplitude is also influenced by the amplitude and duration of the MUAPs (see equation 2 in Farina et al. 2014). Additionally, the estimation of sEMG amplitude is always lower than the sum of individual MUAP amplitudes due to a phenomenon known as amplitude cancellation (Day and Hulliger 2001; Keenan et al. 2005; Negro et al. 2015). In addition, factors such as muscle fiber properties (e.g., MUAP size and conduction velocity) (Farina et al. 2014; Del Vecchio et al. 2017), muscle architecture (Vieira et al. 2017), and inter-electrode distance and placement (Vieira et al. 2023) have been shown to directly affect sEMG amplitude. Therefore, the higher activation in the distal region of the biceps femoris long head during knee flexion cannot be necessarily inferred as an increased neural drive to this region compared to the proximal region, as differences in sEMG amplitudes between anatomical locations may also be influenced by MUAP amplitude (Martinez-Valdes et al. 2018). To better address this issue, we decomposed HDsEMG signals to analyze individual motor unit activity, allowing us to examine both central and action potential properties and provide more accurate insights into the mechanisms underlying the observed activation differences.

### Motor unit action potential properties, not discharge rate modulation, explain localized activation in the biceps femoris long head

A possible explanation for the regional activation within a muscle is the differential modulation of presynaptic inputs to alpha motor neurons that innervate specific regions (Watanabe et al. 2021), as suggested in muscles such as the first dorsal interosseous during flexion and abduction tasks (Desmedt & Godaux 1981) and the rectus femoris during cutaneous electrical stimulation (Watanabe et al. 2015). The most reliable method to investigate this possibility is through the assessment of individual motor unit central properties (mean discharge rate, discharge rate variability and CoV-ISI) as well as time- and frequency-domain coupling between motor unit discharge patterns, as conducted in several recent studies (Cabral et al. 2018; Cabral et al., 2024; Cohen et al. 2020; Sahinis et al. 2024; Crouzier et al. 2024). Regarding the hamstrings, recently, Sahinis et al. (2024) observed distinct neural drive along the semitendinosus muscle during isometric knee flexion tasks. Specifically, the mean discharge rate was greater for motor units in the proximal region compared to the distal region. Moreover, neural drive variability was higher in the proximal at the three lower target forces (10%, 20%, and 40%) and across all three knee joint angles (0°: long length, 45°: intermediate length, and 90°: short length). These findings suggest region-specific motor unit discharge characteristics and neural control in the semitendinosus across varying muscle lengths.

In contrast, our study did not observe any difference in mean discharge rate (Figure 4), CoV-ISI values, or recruitment and de-recruitment thresholds (**Online Resources 1 and 2**) between the proximal and distal regions of the biceps femoris long head. These results indicate that, unlike the semitendinosus, motor units located in different proximodistal regions of the biceps femoris long head are modulated similarly during isometric tasks. Given that both of these hamstring muscles are biarticular and share similar functions (Kellis and Blazevich 2022), it is intriguing that motor units in the semitendinosus exhibit region-specific neural drive during knee flexion, while those in the biceps femoris long head do not. There are several potential explanations for these differences. Although both muscles are innervated proximodistally by different branches of the sciatic nerve (Seidel et al. 1996), the semitendinosus has two distinct proximodistal regions separated by a tendinous inscription, with each region innervated by a separate nerve branch (see Figure 2 in Woodley & Mercer, 2005). This specific neuroanatomical organization may facilitate region-specific motor unit modulation in this muscle, as observed by Sahinis et al. (2024). Additionally, the presence of the biceps femoris short head may contribute to the differences between semitendinosus and biceps femoris long head. For instance, the short head is innervated separately from the long head by a single nerve branch originating from the common fibular nerve (Farfán et al. 2024). This distinct innervation arrangement could influence the modulation of neural drive along the long head, potentially resulting in less pronounced proximodistal motor unit activity during both knee flexion and hip extension tasks. Although these potential explanations provide insight into the observed differences, further research is required to explore motor unit discharge properties across different regions of both the semitendinosus and biceps femoris long head, ideally through simultaneous recordings.

To investigate whether potential proximodistal differences in the MUAP properties of biceps femoris long head motor units could explain the observed differences in sEMG amplitude, we calculated MUAP amplitude and MUAP conduction velocity, which are two commonly used parameters in the literature (Martinez-Valdes et al. 2018; Valli et al. 2024). Consistent with the sEMG amplitude results, higher MUAP amplitudes and conduction velocities were observed in the distal region of the biceps femoris long head compared to the proximal region. Changes in these motor unit action potential properties, rather than central properties, have previously been associated with changes in sEMG amplitude (Martinez-Valdes et al. 2018). Collectively, these findings suggest that the greater sEMG amplitudes observed in the biceps femoris long head during knee flexion and hip extension tasks are primarily driven by action potential properties, rather than discharge statistics.

The localized differences in motor unit action potential properties, and consequently bipolar sEMG amplitude, in the biceps femoris long head are likely explained by anatomical and mechanical differences within the muscle. Miyamoto & Hirata (2021) recently reported that during isometric knee flexion at 20% of MVC, the distal region of the biceps femoris long head exhibited greater stiffness, as measured by supersonic shear wave elastography, compared to the central and proximal regions. The authors also observed higher sEMG amplitudes in the distal region, which aligns with both the current study and previous findings (Schoenfeld et al. 2015; Hegyi et al. 2018, 2019). This increased stiffness in the distal region may result from architectural variations within the muscle, as the biceps femoris long head has longer fibers and a greater pennation angle proximally, which could lead to reduced strain in the proximal region (Kellis 2018). While this relationship between localized peripheral motor unit activity, sEMG amplitude and the muscle’s architecture appears plausible, further investigation is required to fully understand these mechanisms.

### Methodological considerations and limitations

When assessing muscle activation using conventional bipolar electromyography, several critical factors must be considered such as the location of the innervation zone (Mancebo et al. 2019), the direction of the task (Lulic-Kuryllo et al. 2022), and the selection of representative sites within muscle regions (Cabral et al. 2022; Albarello et al. 2022). Our results highlight such considerations, suggesting that bipolar sEMG amplitudes obtained from a single region of the biceps femoris may not accurately represent global muscle activity during knee flexion and hip extension tasks, likely due to peripheral factors.

However, some limitations of our study should be acknowledged. First, the relatively low number of motor units identified in each region may limit the generalizability of our findings. Future research should aim to address this by incorporating larger and denser surface grids of electrodes, which could potentially capture a greater number of motor units (Caillet et al. 2023). Second, we assessed the biceps femoris long head at a single muscle length during both knee flexion and hip extension tasks. Considering recent evidence suggesting that changes in muscle length directly influence the shared synaptic oscillations to spinal motor neurons (Cabral et al. 2024), it would be interesting to investigate whether similar responses occur when altering knee and hip joint angles. Additionally, while the proximal grid of electrodes was positioned as close as possible to the origin of the biceps femoris (see *Methods*), the superficial nature of the gluteus muscle compromised optimal placement. In this regard, techniques such as functional magnetic resonance imaging could enhance precision in analyzing the proximal region of the biceps femoris (Schuermans et al. 2014). Lastly, due to the limited number of probes available, the activity of other hamstring muscles was not explored in this study, which represents another avenue for future research.

## Conclusions

Considering that the long head of the biceps femoris shows the highest prevalence of injuries among the hamstrings, with injuries occurring in the proximal, central, and distal regions of this muscle (Grange et al. 2023), this study provides interesting insights into the proximodistal differences in biceps femoris long head activity during isometric knee flexion and hip extension tasks. Our findings indicate that greater sEMG amplitude in the distal region of the biceps femoris long head cannot be attributed to differences in neural drive between proximal and distal regions, as no significant variations were observed in central motor unit properties. Instead, the increased distal activation appears to be driven by motor unit action potential properties, likely influenced by intra-muscular anatomic and mechanical variations. Furthermore, this study emphasizes the limitations of relying solely on sEMG amplitude to infer neural control strategies across different regions of the biceps femoris long head.

## Supporting information

Online Resource 1

Online Resource 2

## Acknowledgements

This study was funded by CAPES, FAPERJ and FINEP. We would like to thank Dr. Taian Vieira for his valuable feedback on the manuscript, which helped us refine the terminology and enhance the interpretation of our findings.

## Authors’ Contributions

All the authors contributed substantially to the manuscript. Conception and design of the experiments (JCSA, HVC, LFO), collection of data (JCSA), analysis of data (JCSA, HVC, FN), interpretation of data (JCSA, HVC, FN, LFO), drafting the article and revising it critically for important intellectual content (JCSA, HVC, FN, LFO), and final approval of the version to be published (JCSA, HVC, FN, LFO). All authors have read and approved the final version of the manuscript and agree with the order of presentation of the authors.

