## Supplementary figures and images for "Greater distal activation of the biceps femoris long head reflects proximodistal differences in motor unit action potential properties"

### Online Resource 1

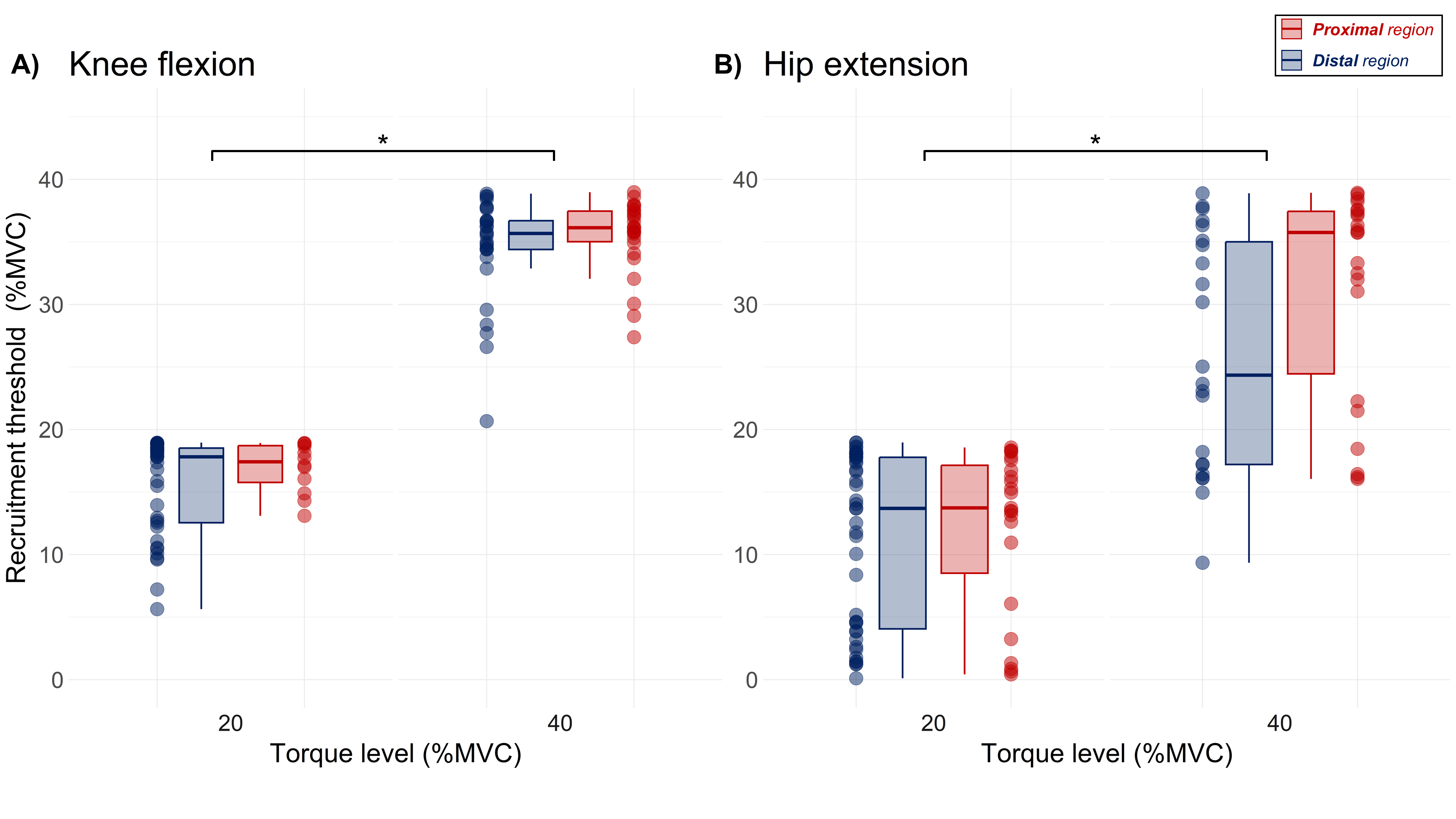

### Online Resource 2

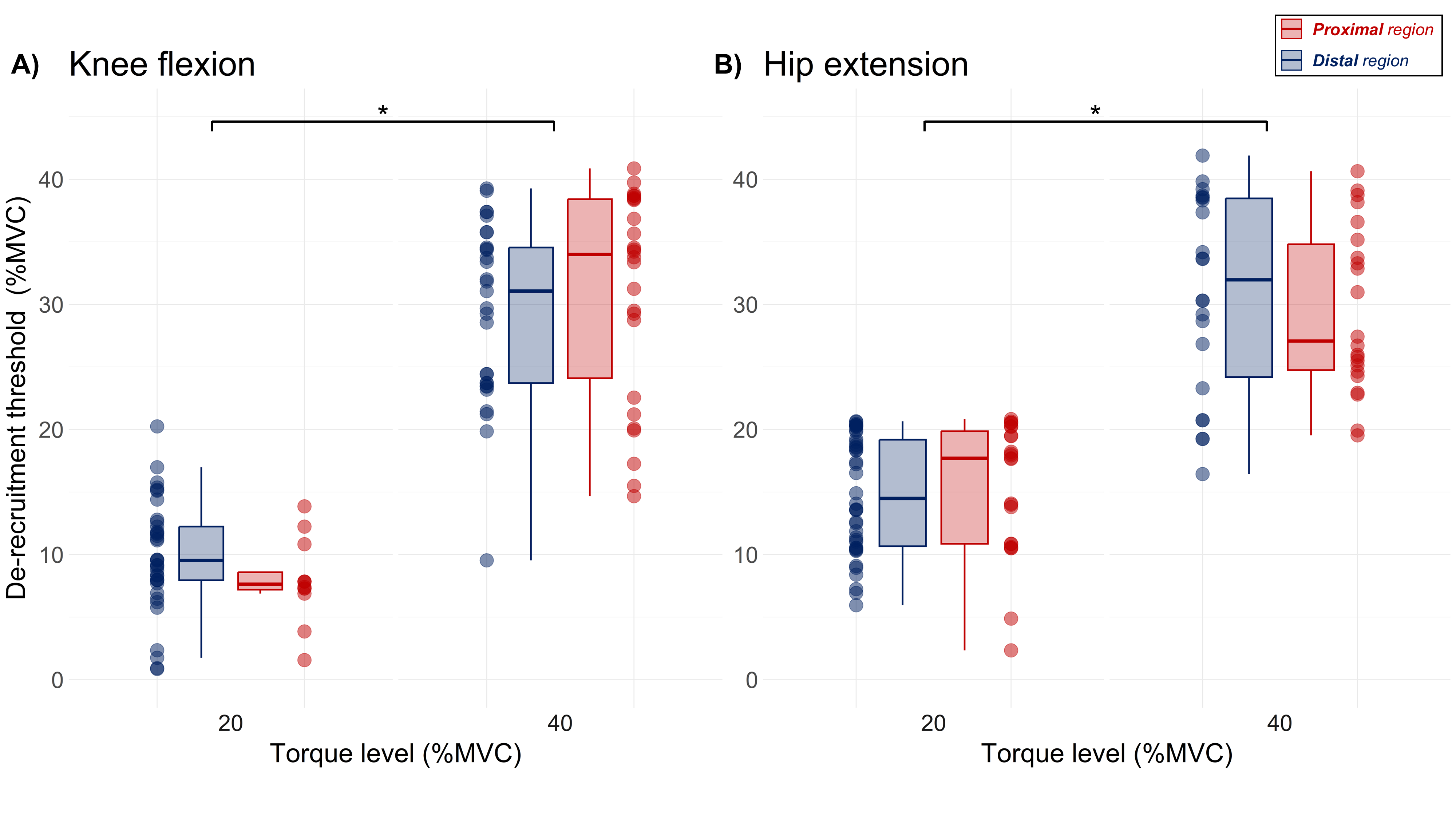
